# Core–periphery dynamics in a plant–pollinator network

**DOI:** 10.1101/543637

**Authors:** Vincent Miele, Rodrigo Ramos-Jiliberto, Diego P. Vázquez

## Abstract

Mutualistic networks are highly dynamic, characterized by high temporal turnover of species and interactions. Yet, we have a limited understanding of how the internal structure of these networks and the roles species play in them vary through time. We used six years of observation data and a novel statistical method (dynamic stochastic block models) to assess how network structure and species’ structural position within the network change across time in a quantitative plant–pollinator network from a dryland ecosystem in Argentina. Our analyses revealed a core–periphery structure persistent through seasons and years. Yet, species structural position as core or peripheral were highly dynamic: virtually all species that were at the core in some seasons were also peripheral in other seasons, while many other species remained always peripheral. Our results illuminate our understanding of the dynamics of ecological networks and have important implications for ecosystem management and conservation.

## Introduction

Studies of plant–animal mutualisms have historically focused on the interactions between one or a few plant species and their animal mutualists (1; 2). This approach guided decades of research, illuminating our understanding of the natural history, ecology and evolution of plant–animal mutualisms, but at the same time limiting our understanding of how interactions operate in their broader community context (3). More recently, the use of a network approach to study of plant–animal mutualistic interactions in their community context has offered new insights on the relative specialization and reciprocal dependence of these interactions and, ultimately, the ecological and evolutionary processes that depend on them (4; 3; 5; 6). The study of mutualistic networks has revealed several pervasive properties, including nestedness (7), modularity (8), and asymmetry in both specialization (9) and interaction strength (10), all of which are believed to have important ecological and evolutionary implications (11; 12; 6; 13).

Mutualistic networks are also characterized by high temporal variability, with species and interactions switching on and off through time. In other words, these networks exhibit high temporal turnover of species and interactions (14; 15; 16), in spite of an apparent stability in some aggregate network attributes such as connectance and nestedness (14; 17). Past studies have shown that the most persistent interactions are those located at the network core (the most densely connected region of the network), which usually involves abundant, frequently interacting species, and many occasional peripheral species (16; 18). What we still don’t know is the extent to which the structural position of individual species as core or peripheral varies through time. In other words, is there a persistent set of core species that form the backbone of the network over seasons and years? Or is the core itself also highly dynamic, with species switching between core and peripheral positions?

Answering the above questions is essential to improve our understanding of how different species contribute to community stability and to guide management and conservation efforts. For example, the existence of a stable set of species at the network core could represent a reasonable target for biodiversity conservation—a small, manageable set of keystone species on which to focus conservation efforts (19; 20; 21; 22; 23). Conversely, a highly dynamic network core would make that target more elusive, with a larger, variable set of potentially keystone species.

Here we evaluate how the structure of a plant–pollinator network and the structural position of species in the network change across time. We focus on a previously published bipartite, weighted (non-binary) plant–pollinator network spanning six years in a dryland ecosystem in Villavicencio Nature Reserve, Argentina (16). Our network representation focuses on the relative ecological effects between pairs of interacting species (usually referred to as *dependences*, 10; 24). Using a recent statistical framework (dynamic stochastic block models, hereafter dynSBM) (25), we quantify the temporal switching of the structural position of plants and pollinators. This analysis allows us to provide a comprehensive picture of the temporal dynamics of the internal structure of this mutualistic network.

## Material and methods

### Study site and data collection

We used a dataset describing a plant–pollinator network from pollinator visits to flowers in a dryland ecosystem. Data were collected weekly during three months during the flowering season (Austral spring and early summer, September–December) between 2006 and 2011 from the Monte Desert ecoregion at Villavicencio Nature Reserve, Mendoza, Argentina (32° 32’ S, 68° 57’ W, 1270 m above sea level). The data include 59 plant species, 196 flower visitor species, and 28015 interaction events (flower visits) involving 1050 different pairs of interacting species. Plant abundance was estimated based on the density of flowers of each plant species, as flowers are the relevant plant structure for this interaction type. Flower abundance was estimated during the flowering season of all study years using fixed quadradts/transects. Several rare plant species were absent from our fixed quadrats and transects but present elsewhere in our study site; for those species we assigned an abundance of one flower, the minimum we could have detected with our sampling method. A full account of the methodology can be found in Ref. (16; 22).

### Building plant–pollinator dependence networks

We aggregated the data by pooling the number of visits of any pollinator to any plant in 3 subseasons by year (before November 1st, after November 30th and in between). Such level of aggregation allowed us to consider seasonal dynamics at a temporal grain that was not too fine nor too coarse to allow a reasonable representation of network structure.

For any subseason, we built a plant-pollinator *dependence network D*, a directed weighted network representing the relative dependences among plant and pollinator species (10; 24). From the number of visits in a time interval *X_ij_* between any pair of species of plant and pollinator (*i,j*), we considered two directed and weighted edges in *D*: the dependence of plant *i* on pollinator *j*, 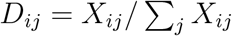, representing the number of visits of pollinator *j* to plant *i* divided by the total number of visits received by plant *i*; and the reciprocal dependence of pollinator *j* on plant *i*, 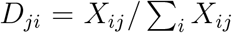, representing the number of visits of pollinator *j* to plant *i* divided by the total number of visits done by *j*. Applying this approach to our raw data, we obtained a time series of 18 dependence networks. To represent graphically these networks, we showed the successive bi-adjacency matrices (plants in rows, pollinators in columns) using a color code accounting for the two values *D_ij_* and *D_ji_* for any species pair (*i, j*) (see an example in Figure 1).

**Figure 1:**
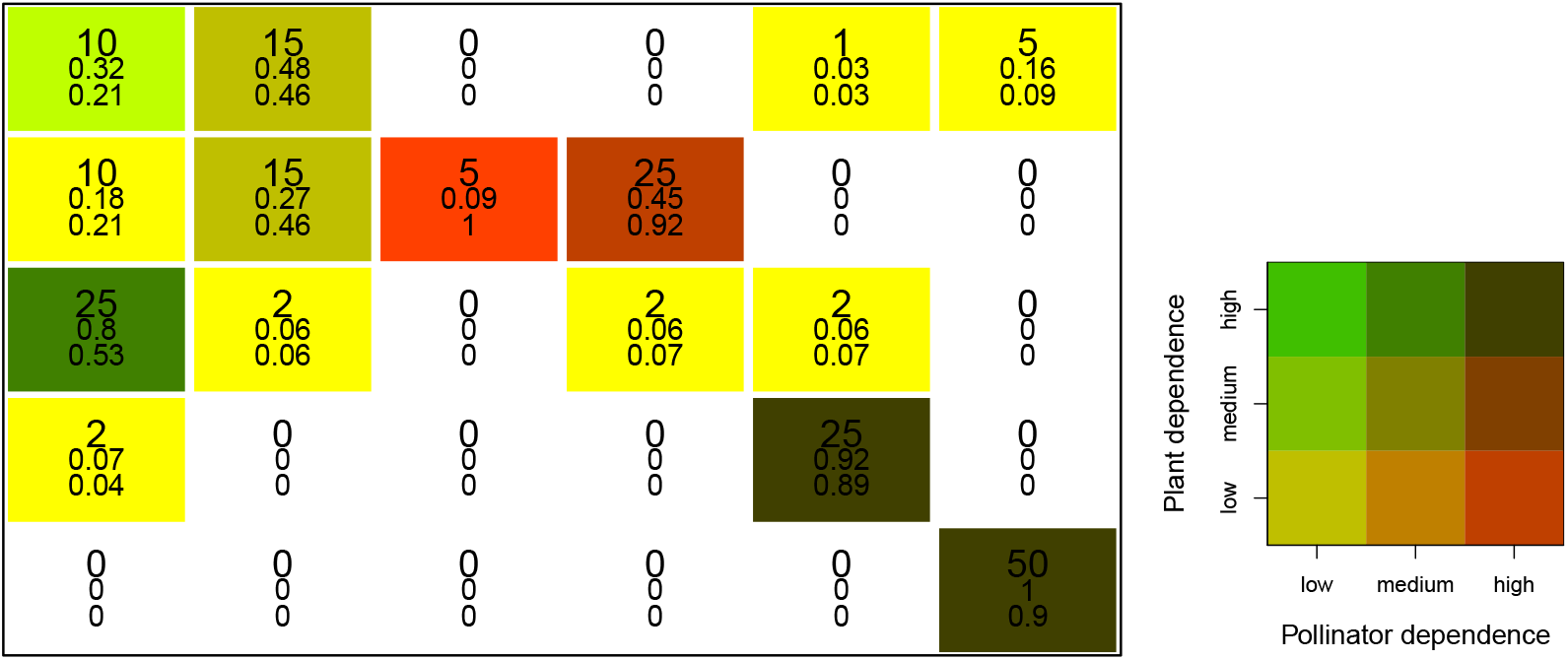
Representation of a dependence network between 5 plant species (rows) and 6 pollinator species (columns). Each cell is colored according to the legend and filled with the number of visits (top value in larger font), the plant and pollinator dependence values *D_ij_* (middle) and *D_ji_* (bottom). The legend shows the color code accounting for the two dependence values for any species pair (*i,j*) (darker green represents higher the plant dependence; stronger red represents higher the pollinator dependence). This example shows the advantage of studying dependence values instead of raw data. The number of visits in cells (3,1), (2,4) and (4,5) are all equal to 25. Meanwhile, these number of visits do not characterize the same kind of interaction, as shown by the dependence values. Indeed, plant 3 is highly dependent on pollinator 1 (the reverse is not true), pollinator 4 is highly dependent on plant 2 (the reverse is not true) whereas plant 4 and pollinator 5 are mutually dependent and have a quasi-exclusive relationship. Lastly, the number of visits in cell (5,6) is twice the number in cell (4,5) but the dependence values are comparable (dependence is scale invariant).

### Inferring topology and species’ structural position in the dynamic network

Recently in Ecology (26; 27; 28; 29; 30), some authors have suggested the use of statistical methods which jointly infer structural properties and species positions. Originally developed in the field of social sciences (31), *Stochastic Block Models* (SBM; 32; 33)—also called *Group Models* in the seminal work by Allesina and Pascual (27)—aim at grouping nodes (species in our case) that are statistically equivalent, “acting” similarly in the network, *i.e*., having an equivalent “structural position”. These methods follow a particular paradigm: instead of searching for a particular pattern, we infer one from the data. SBM can handle weighted networks with appropriate statistical distributions; we chose them for their ability to decipher core-periphery structure in network data (as mentioned in Figure 1 in 34), as they can infer groups of core species and peripheral species.

Furthermore, studying network dynamics requires a method that can handle and model the whole time series of network snapshots (i.e., in a *dynamic network*). Recently, Matias and Miele (25) proposed an extension of SBM for dynamic networks called dynSBM. Under this approach, the structural position of any species can vary over time. In other words, each structural group (for instance a core group) is inferred using the complete series, but the group membership can vary from any time step to another. Here we rely on a modified version of this approach dedicated to bipartite networks (see Supplementary information) implemented in the R package `dynsbm` available on CRAN at https://cran.r-project.org/web/packages/dynsbm/. Importantly, the number of groups is constant and selected with an appropriate heuristics (Supplementary Figure S1).

## Results

### A persistent core-periphery structure

By applying the dynSBM algorithm, we found that the Villavicencio plant–pollinator network is organized as a core–periphery structure. This network structure comprises two components, each one composed of a group of plants and a group of pollinators. The first component consists of one group of plant species and one group of pollinator species forming a persistent cohesive module (the network *core*), while the second component was composed of a group of plants and a group of pollinators gravitating in the network *periphery* (Supplementary Fig. S1). The proportions of species in these groups varied only modestly through time (*χ*^2^ = 64.92, d.f. =51, *P* = 0.09) in spite of being unconstrained in dynSBM (Fig. 2); in contrast, these proportions varied widely in randomized networks (Supplementary Figure S3).

**Figure 2:**
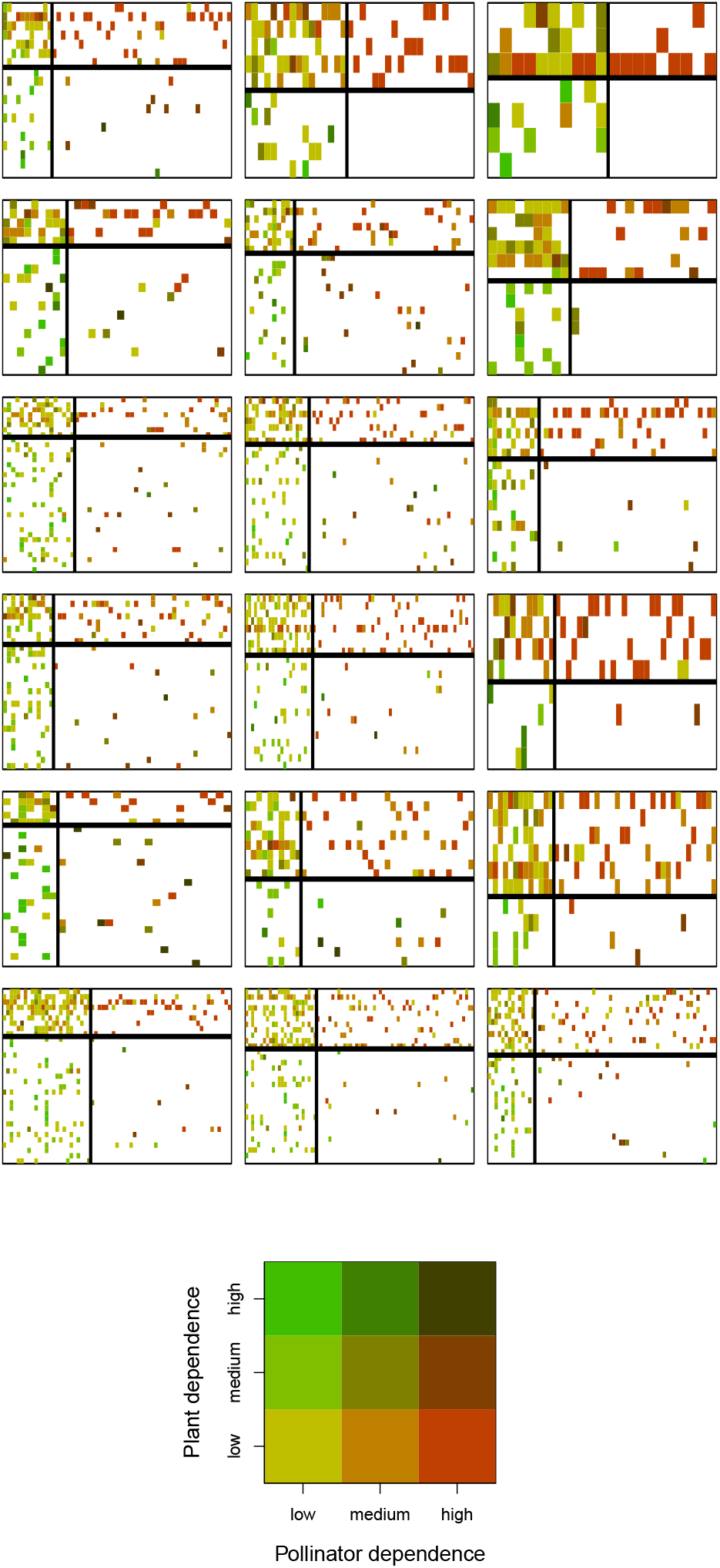
Temporal dynamics of Villavicencio plant–pollinator network. For each matrix, cells represent the plant and pollinator dependence values between a plant (rows) and pollinator (columns) species, with a color computed as a mixture of the two dependence values according to the legend. Rows and columns were reorganized according to the dynSBM group membership: dark lines separating each matrix delineate the group boundaries (core/peripheral group of plants above/below the horizontal line; core/peripheral group of pollinators on the left/right of the vertical line).

Core and peripheral species differ markedly in terms of their linkage patterns. The core group of plants (top rows of matrices in Fig. 2) consisted of species visited by many pollinator species, especially species in the core group of pollinators (left columns of matrices in Fig. 2), which visited many plant species. Species in these core groups of plants and pollinators are weakly dependent on their interaction partners (Supplementary Fig. S2). Thus, the network core can be envisioned as a densely connected “module” of generalized plant and pollinator species with low mutual dependence among them (Fig. 3). In contrast, the peripheral group of plants (bottom rows of matrices in Fig. 2) includes species visited mostly by core pollinator species; dependence is highly asymmetric for these plants, in the sense that they are highly dependent on pollinators who are not reciprocally dependent on their host plants (Supplementary Fig. S2). Likewise, the peripheral group of pollinators (right columns of matrices in Fig. 2) includes species interacting mostly with core plants, also asymmetrically dependent on plants that are not reciprocally dependent on them (Fig. 3). In addition, there are only a few interactions between peripheral plant and pollinator species, with no particular trend regarding their reciprocal dependence (Supplementary Fig. S2).

**Figure 3:**
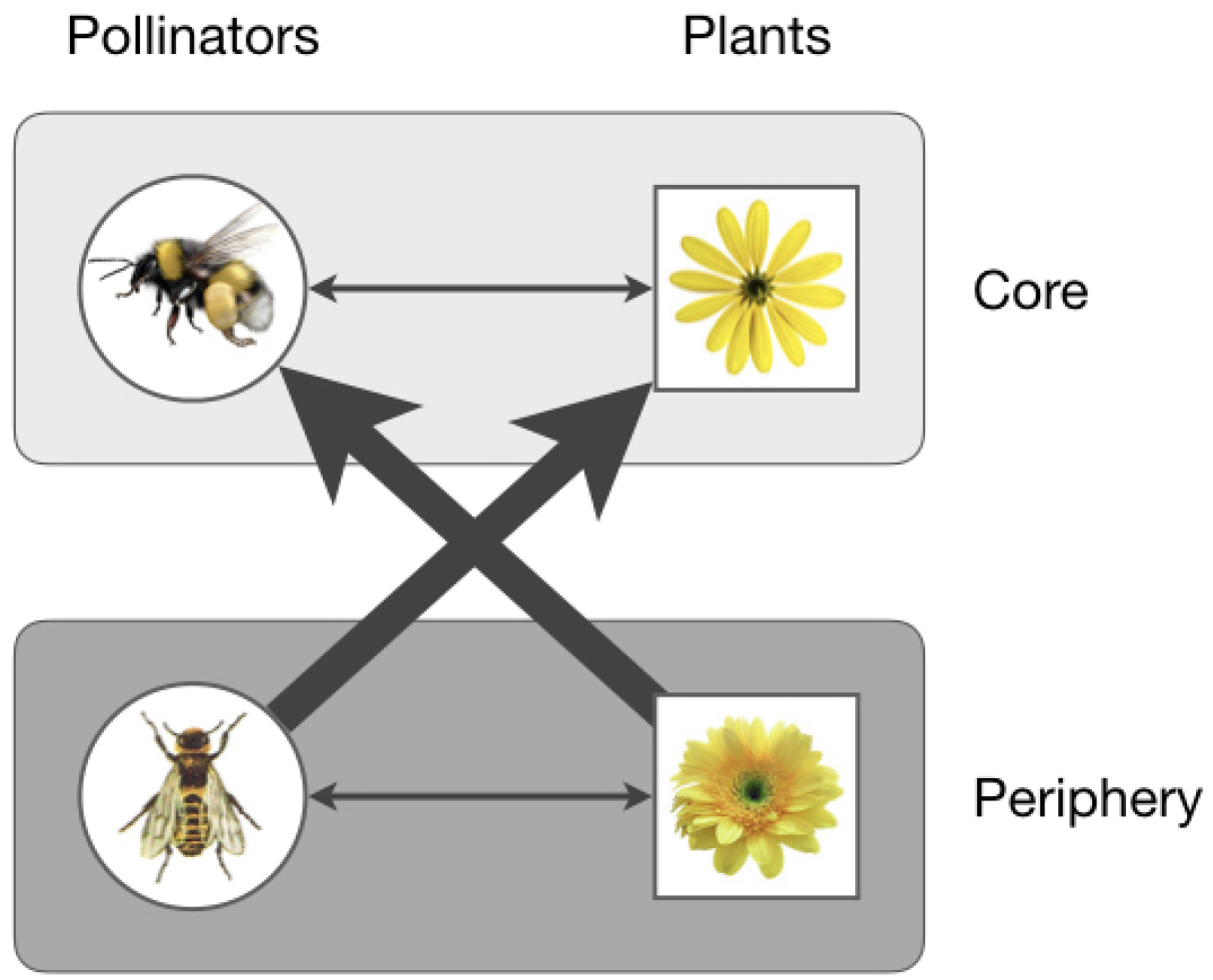
Graphical representation of the core–periphery structure found in our dynamic plant–pollinator network. Arrows depict dependences of one species (arrow origin) on another (arrow tip). Arrow widths are proportional to typical dependence values between groups. Pollinators/plants of the network periphery are strongly dependent on plants/pollinators that belong to the network core.

### The core–periphery structure is robust to changes in species diversity and composition

The core–periphery structure persisted despite two sources of variation: the diversity of species and their identities. First, the diversity of plant and pollinator species varied over time, so that each year the number of plant species in bloom tended to decrease from the first to the third subseason, whereas the number of pollinator species species tended to peak in the second subseason (Supplementary Fig. S4); yet, the proportion of core plant species increased from the first to the third subseason each year (Fig. 2; plant core group in the upper part of each matrix). Thus, the size of the plant core group was independent of plant diversity. Second, the identity of interacting species and their activity (as measured by the total number of floral visits received by a plant or performed by a pollinator) changed greatly from one time step to another, resulting in substantial temporal variation in the species assembly (Supplementary Fig. S5). Yet, despite these variations in the interactions at the species level, the core–periphery structure persisted over time.

### Species in the core are also sometimes peripheral

Species structural positions were highly dynamic. Almost all species that were in the core in some seasons were also peripheral in other seasons (except one plant and one pollinator species); however, a large proportion of peripheral species never became part of the core (52% for plants, 72% for pollinators; see Fig. 4). Thus, only a subset of species were ever part of the core, and virtually no species occupied that position persistently through time.

**Figure 4:**
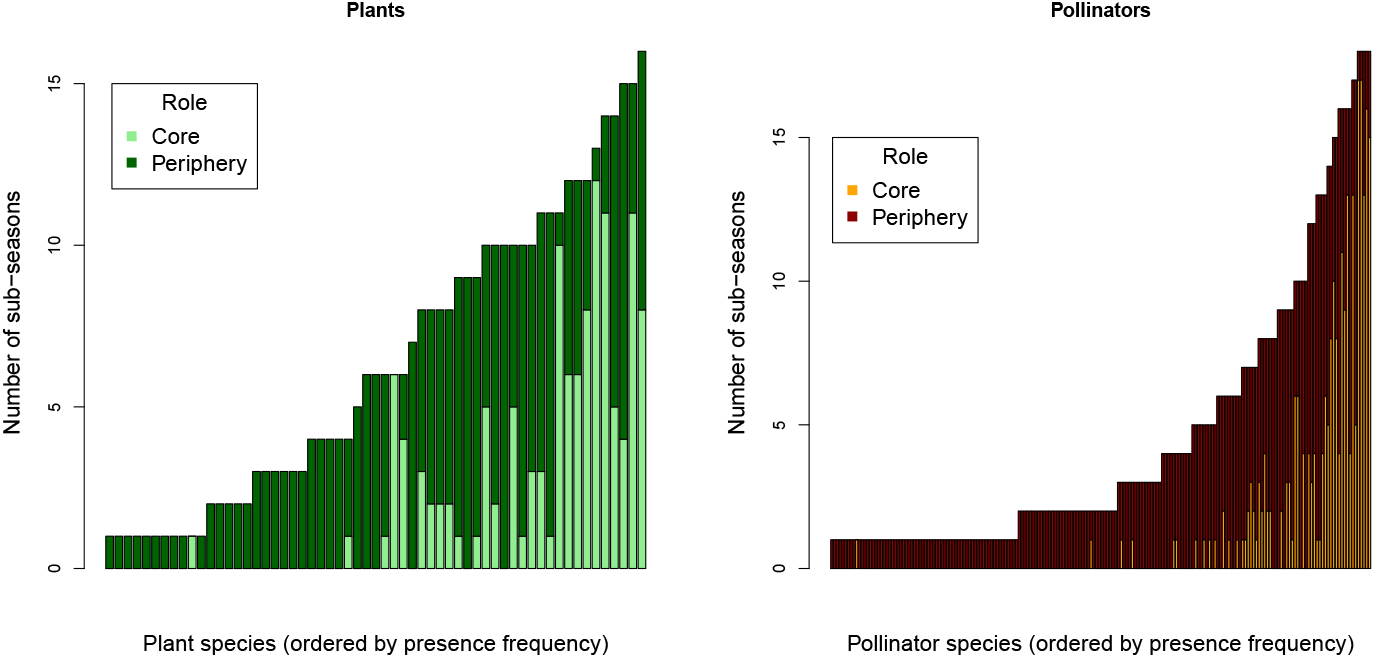
Temporal shifts in species structural positions. Each stacked bar (one by species) represents the number of subseasons any species was found in the core (light color) or in the periphery (dark color). Bars were ranked according to the number of subseasons any species was observed and present in the network. For plants (left) and pollinators (right).

There was a positive correlation between overall species presence (i.e. the number of subseasons a species was recorded interacting) and their presence in the core: the more frequently a plant or a pollinator species was present in the community, the more frequently it was found in the core (see Fig. 4 and Supplementary Fig. S6). Furthermore, for plant species for which we have independent abundance data, we observed that their abundance tended to be higher when they are in the core than when they are peripheral (Supplementary Fig. S7).

## Discussion

Our analysis using dynamic stochastic block models allowed us to delve into the topological dynamics of a plant-pollinator network. In a nutshell, we found that this network is characterized by a core–periphery structure persistent through seasons and years, while exhibiting high temporal switching of species structural positions. These results offer a unique temporal perspective into the dynamics of mutualistic networks.

The core–periphery structure was maintained in spite of high temporal variation in species richness and composition. The distribution of dependences also persisted over time, with highly asymmetric dependences for most peripheral species, which tended to interact with core species; in turn, interactions among core species tended to be more symmetric, albeit with weaker dependences. Yet, the network position occupied by plant and pollinator species was highly dynamic: virtually all species that played a core role in some seasons were also peripheral in other seasons, while many other species remained always peripheral. Furthermore, presence in the network core was related to overall species presence: species present in many subseasons tended to be more consistently at the core than species present only in few subseasons. Previous studies had documented that nestedness (which can be viewed as a particular type of core–periphery structure, 35) characterizes many plant–animal mutualistic networks (7) and that such structure is persistent over the years (14; 16) in spite of an enormous temporal variation in the occurrence of interactions (14; 15; 16). Our findings extend those results, indicating that species structural position in the network is also highly dynamic. Thus, while the core–periphery structure persists over time, the taxonomic identity of the core changes drastically through seasons and years, and no species can be identified as playing permanently a core role.

The latter finding has far-reaching practical implications, as the idea of focusing management and conservation efforts on a small subset of species at the network core (19; 20; 21; 22; 23; 36; 37) may be difficult to achieve, given that virtually no species plays that role consistently over time in the long run. Our findings do indicate that a small subset of species is likely to be found playing a key role as part of the network core in many seasons and years, which brings them close to the notion of “core” species and would make them adequate targets for conservation efforts. Plant species in this group include *Condalia microphilla, Larrea divaricata, Prosopis flexuosa* and *Zuccagnia punctata* whereas flower visitors in this group include *Apis mellifera, Augchloropsis* sp., *Bombus opiphex, Centris brethesi, Copestylum aricia*, and *Xylocopa atamisquensis*.

Yet, a majority of core species was core in a substantially smaller fraction of subseasons (see Fig. 4). These species could be viewed as *quasi-core* species, in the sense that they are present in the core only intermittently. Thus, the identification of core species based on one or a few years of sampling—as done in most studies published so far—could be misleading, and a single static characterization of an ecological network will fail to reveal its true core–periphery structure. In this sense, the idea of species “coreness” (35; 38) is not just a black-or-white property determined only by the position of a species in a static or aggregated network, but a relative concept determined by the temporal consistency of the position occupied by a species. Therefore, identifying core species as candidates for management actions requires allocating a greater sampling effort into capturing the temporal dynamics of ecosystems, even if this practice implies relaxing efforts to capture some details of community structure and the detection of very rare species, which are unlikely to be part of the network core and to contribute significantly to community robustness to environmental perturbations.

To conclude, we believe these results illuminate our understanding of the dynamics of ecological networks, indicating the persistence of a core–periphery structure in spite of substantial changes in species richness, composition, interactions and structural position in the network. Yet, we believe we have only scratched the surface of the temporal dynamics of ecological networks. One possible avenue for future research would be to apply the methods used here to analyze other datasets, to assess the generality of our findings.

## Acknowledgements

VM thanks Sebastien Ibanez and Hugo Fort for their useful comments. Funding was provided by the French National Center for Scientific Research (CNRS) and the French National Research Agency (ANR) grant ANR-18-CE02-0010-01 EcoNet (VM), CONICYT/FONDECYT grants 1150348 and 1190173 (RRJ), a FONCYT grant PICT-2014-3168 (DPV), the People Programme (Marie Curie Actions) of the European Union’s Seventh Framework Programme (FP7/2007-2013, REA grant agreement 609305) (DPV), and a Bessel Research Award from Alexander von Humboldt Foundation (DPV).

## Author’s contributions

All authors conceived the study and wrote the manuscript. VM conducted the analyses.

## Details of the dynamic stocastic block model analysis

Under the dynamic stochastic block model (dynSBM) approach (which is presented in details in (1) and (2)), structural position assignment is defined not only by a SBM (one per time step) but also by a Markov chain that models the switches at each time interval. Here, we rely on a modified version of this approach for bipartite networks where each SBM has the same parameters values at each time step. Each SBM is parametrized by an appropriate statistical distribution. Here we used dynSBM with multinomial distributions to model edge weights (dependence values) that were categorized into three levels corresponding to low, *medium* and *high* dependence (lower than 0.2, in between and larger than 0.8, respectively). The number of groups is constant and selected with an appropriate heuristics (Supplementary Figure S1). Structural position assignment (i.e., SBM group membership) can change over time, but there is no constraint for the found structure to be present at each time step (see Supplementary Figure S3). This approach can be reproduced with the R package `dynsbm` available on CRAN at https://cran.r-project.org/web/packages/dynsbm/.

### Choice of the number of groups in dynSBM

**Figure S1:**
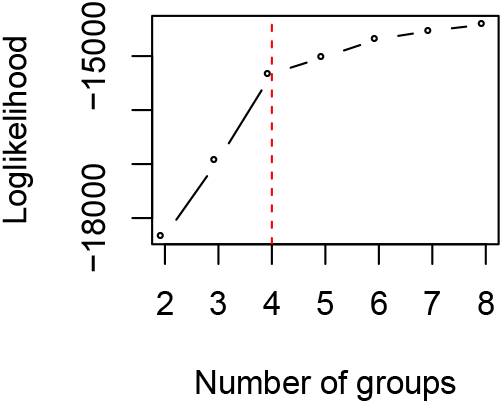
Selecting a dynSBM with 4 groups. The slope of the log-likelihood highly decreases for ≥ 4 groups (“elbow” method, see 2).

### Inter/intra-group dependence values

**Figure S2:**
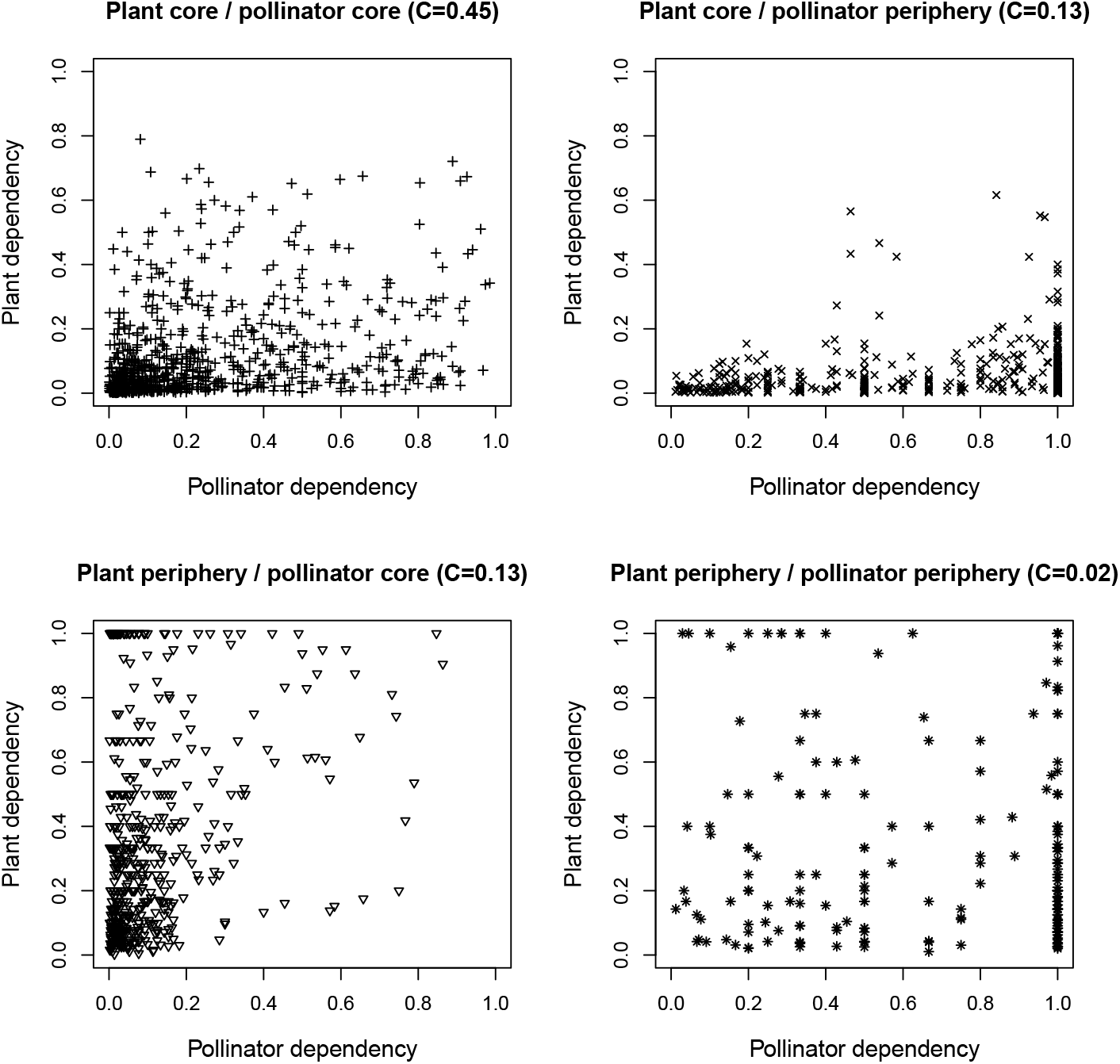
Plant dependence versus pollinator dependence for all the interactions observed in the 18 dependence networks, represented for the four inter-group categories: between plant and pollinator species from the core (top left), from the plant core and the pollinator periphery (top right), from the plant periphery and the pollinator core (bottom left) and from the periphery only (bottom right). The connectance (C) is indicative and corresponds to the fraction of realized interactions.

### No persistence of core/periphery structure in a randomly perturbated network

**Figure S3:**
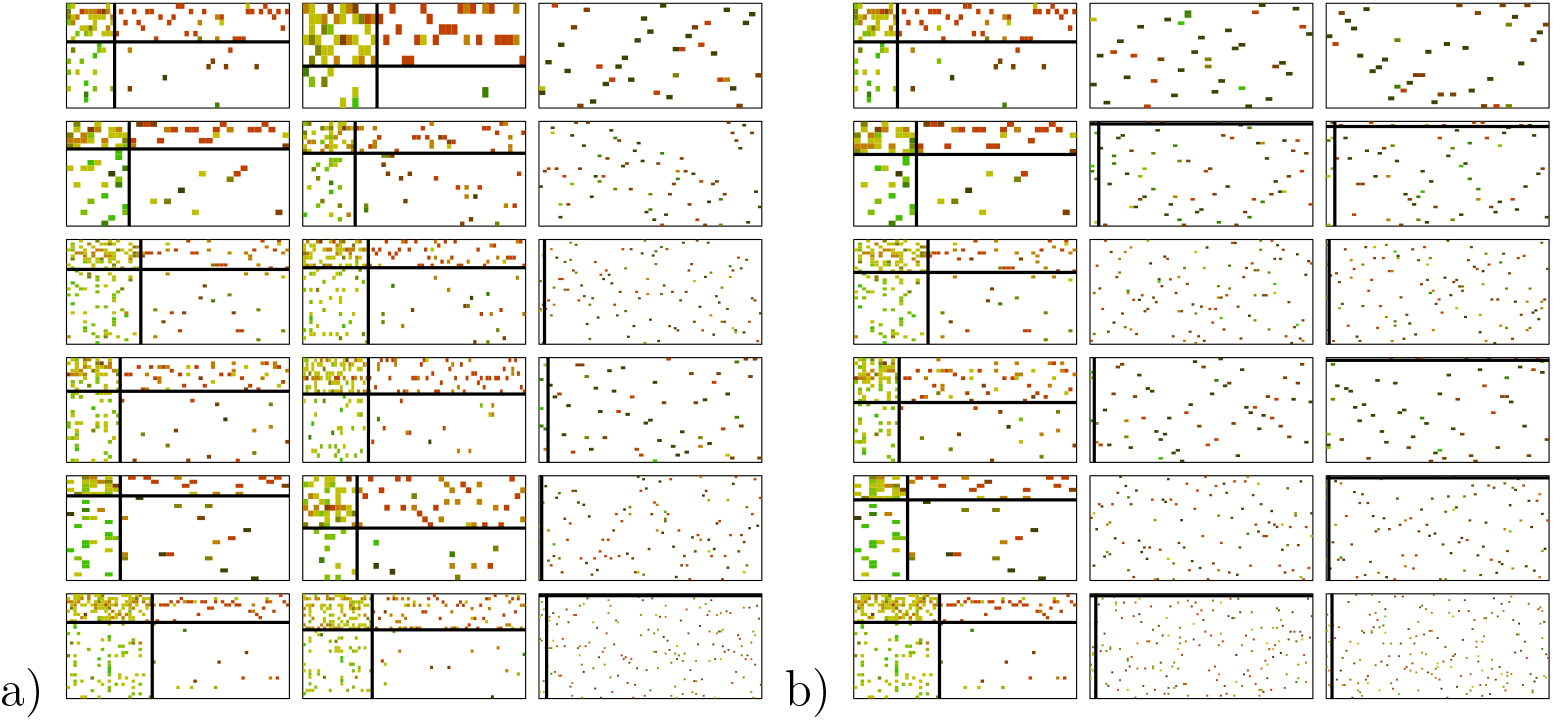
Core/periphery structure does not persist when networks are randomly perturbed. Same as Figure ?? but networks where randomized a) in each third subseasons or b) second and third subseasons. The dynSBM model was re-estimated with a fixed number of 4 groups.

### Variation of the number of species

**Figure S4:**
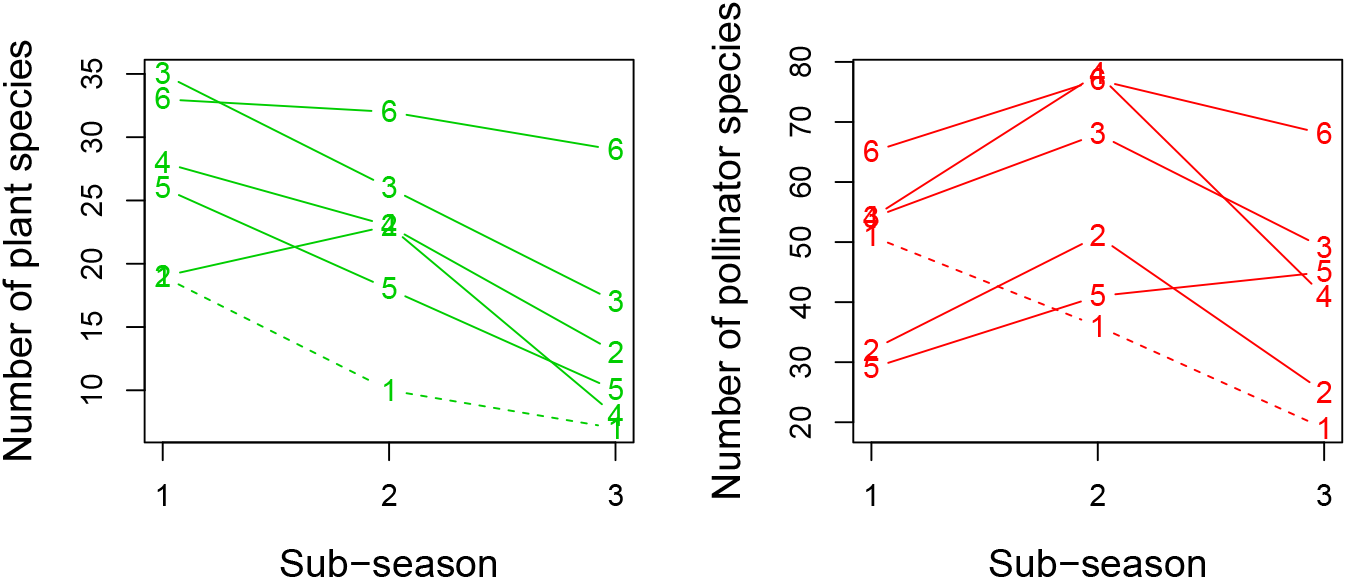
Number of species for the three successive subseasons for each year. The six lines are numbered according to the index of the year of study (from 1 to 6 for 2006 to 2011). The line 1 (2006) is in dashed line to highlight the much lower overall number of species in this particular year. For plants (left) and pollinators (right).

### Temporal variation of species activity

**Figure S5:**
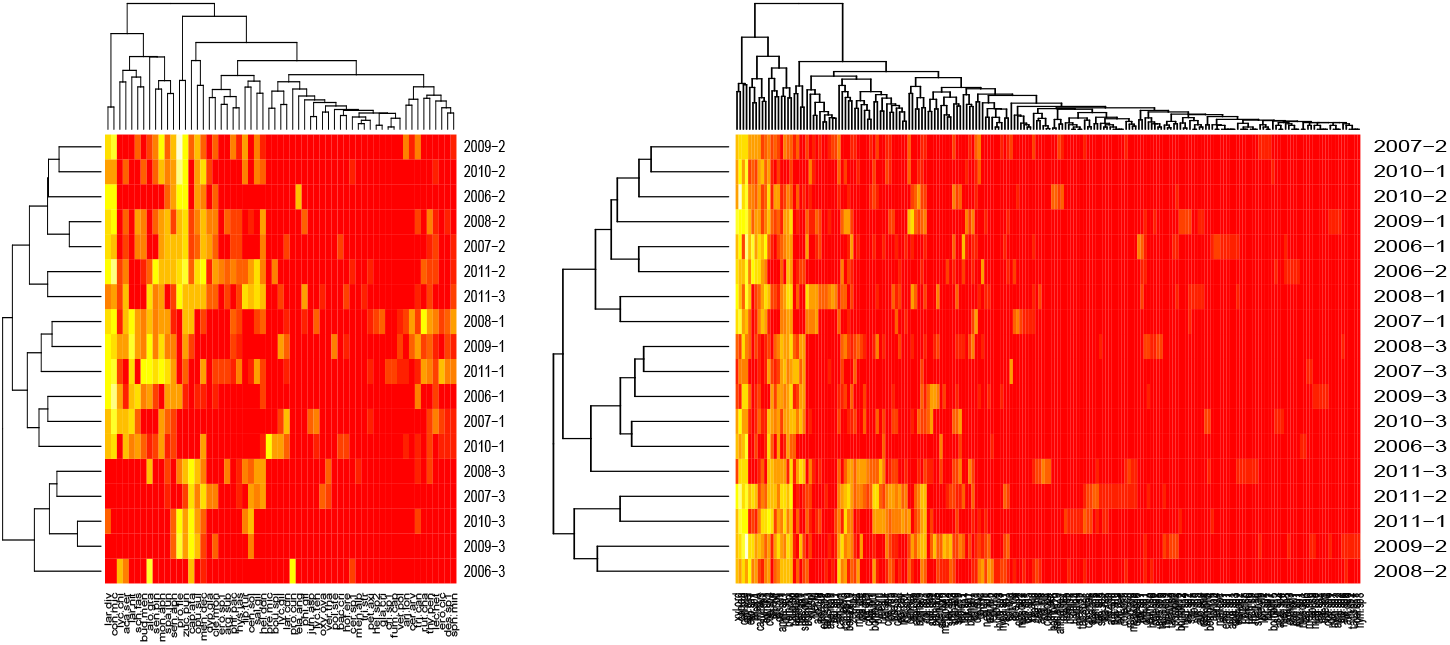
Species activity classification. The activity, measured by the total number of floral visits received by a plant or performed by a pollinator, is represented in a heatmap (red to yellow color scale for null to maximum activity; log-scale). Subseasons in rows (denoted by the year and the index of the subseason in the year) and species in columns are reordered with a hierarchical clustering based on the similarity in activity (log scale, euclidian distance). For plants (left) and pollinators (right). First/second/third subseasons of different years are clearly packed in clumps (*i.e*. subparts of the dendrogram) showing similar plant activity, whereas this pattern is less clear in the case of pollinators (still, the third subseasons are packed together).

### Species turnover and role switch over time

**Figure S6:**
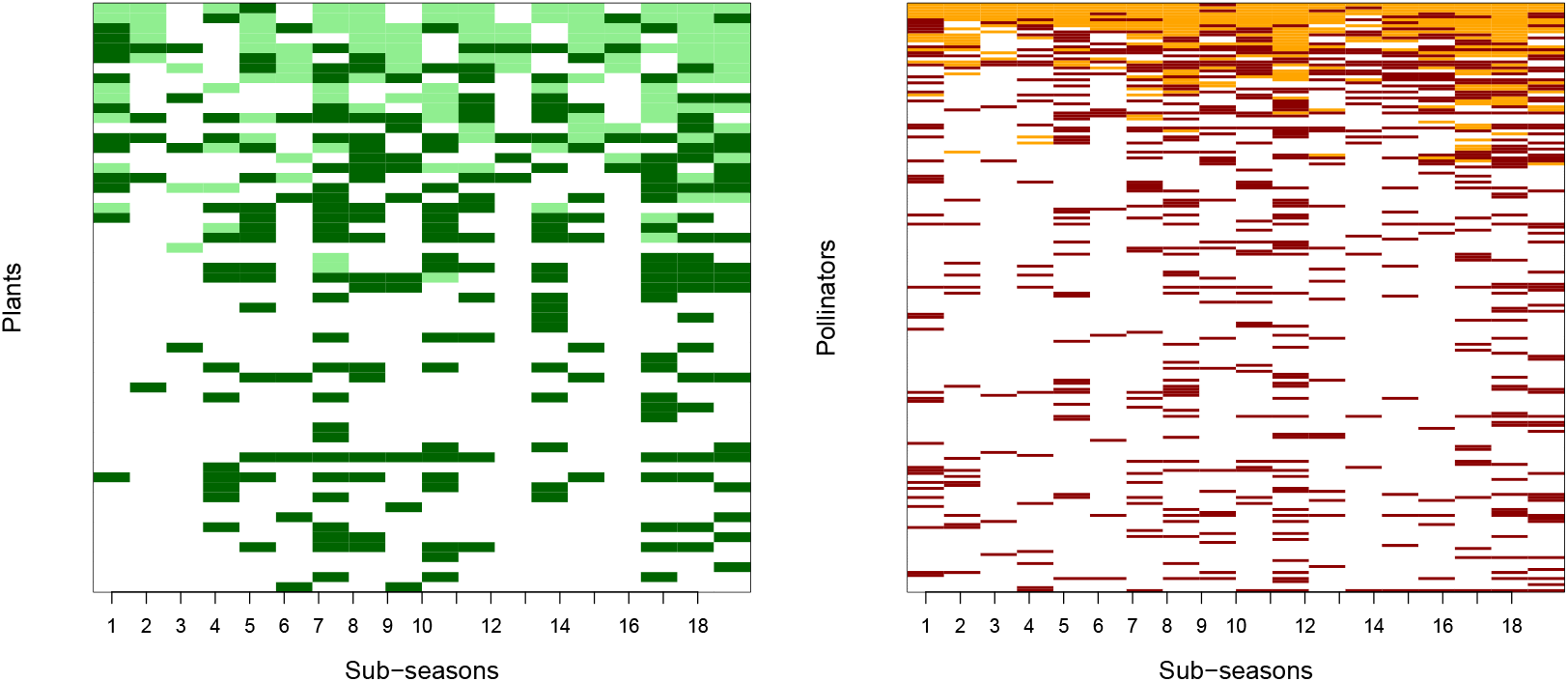
Species turnover and role switch over time. Matrix representation shows when any species (rows) is in the core (light color), in periphery (dark color) or absent (white) over time (columns; 18 subseasons). For plants (left) and pollinators (right).

### Plant role is correlated with abundance

**Figure S7:**
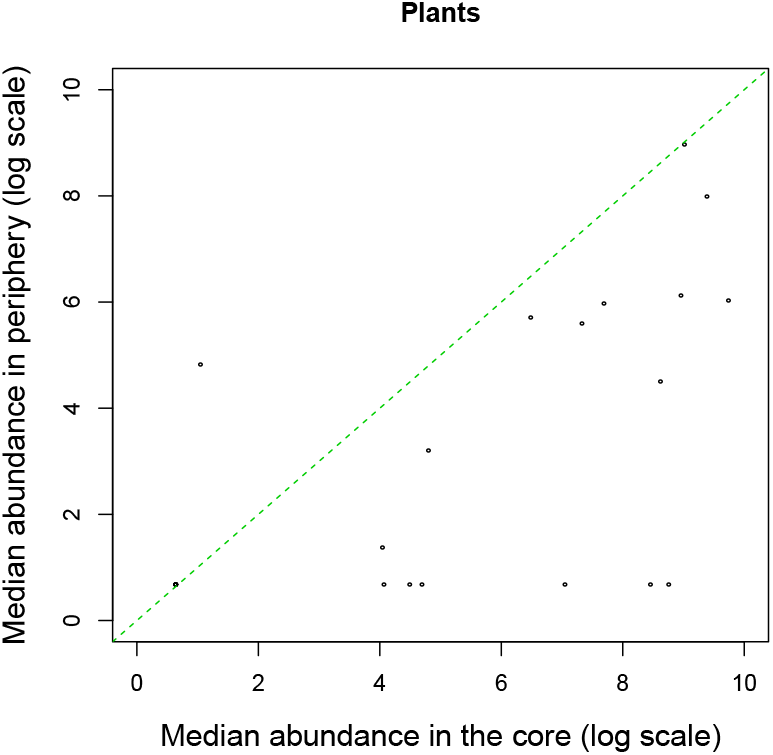
Plant role is correlated with abundance. Each marker shows the median abundance (number of flowers) of a given species when it is in the core compared to its median abundance when it is peripheral. Dashed lines represent the line with slope 1.

